# Insights into the L3 to L4 developmental program through proteomics

**DOI:** 10.1101/2021.05.06.439182

**Authors:** Sasisekhar Bennuru, Zhaojing Meng, James McKerrow, Sara Lustigman, Thomas B Nutman

## Abstract

The establishment of infection with the lymphatic dwelling filarial parasites is dependent on the infectivity and subsequent development of the infective larvae (L3) within the human host to later stages (L4, adults) that require several developmental molts. The molecular mechanisms underlying the developmental processes in parasitic nematodes are not clearly defined. We report the proteomic profiles throughout the entire L3 to L4 molt using an established *in vitro* molting process for the human pathogen *B. malayi*. A total of 3466 proteins of *B. malayi* and 54 from *Wolbachia* were detected at one or more time points. Based on the proteomic profiling, the L3 to L4 molting proteome can be broadly divided into an early, middle and late phase. Enrichment of proteins, protein families and functional categories between each time point or between phases primarily relate to energy metabolism, immune evasion through secreted proteins, protein modification, and extracellular matrix-related processes involved in the development of new cuticle. Comparative analyses with somatic proteomes and transcriptomes highlighted the differential usage of cysteine proteinases (CPLs), BmCPL-1, -4 and -5 in the L3-L4 molt compared to the adults and microfilariae. Inhibition of the CPLs effectively blocked the *in-vitro* L3 to L4 molt. Overall, only 4 *Wolbachia* proteins (Wbm0495, Wbm0793, Wbm0635, and Wbm0786) were detected across all time points and suggest that they play an inconsequential role in the early developmental process.

**Importance:** The neglected tropical diseases of lymphatic filariasis, onchocerciasis (or river blindness), and loiasis are the three major filarial infections of humans that cause long-term disability, impaired childhood growth, reduced reproductive capacity. Global efforts to control and/or eliminate these infections as a public health concern are based on strategies and tools to strengthen the diagnostics, therapeutic and prophylactic measures. A deeper understanding of the genes, proteins and pathways critical for the development of the parasite is needed to help further investigate the mechanisms of parasite establishment and disease progression, because not all the transmitted infective larvae get to develop successfully and establish infections. The significance of this study is in identifying the proteins and the pathways that are needed by the parasite for successful developmental molts, that in turn will allow for investigating targets of therapeutic and prophylactic potential.

## Introduction

Lymphatic filariasis is caused primarily by the parasitic nematodes *Wuchereria bancrofti*, *Brugia malayi* and *Brugia timori*. Infection is initiated when infective L3 larvae enter the human host through the skin and subsequently develop into L4 after a developmental molt. Given that the L3 to L4 molting process occurs across all nematodes and that it is the essential first step towards establishing infection, the molting process has been postulated to be a target for intervention strategies (1). Despite the notion that these early (L3 and L4) developmental stages would be an important target for prophylactic vaccines, the biology of these early mammalian stages of the lymph-dwelling filarial parasites have been not well-studied.

There is evidence that molting and ecdysis in most nematodes are under the control of neurosecretory and endocrine processes (2). While functional nuclear hormone receptors (3, 4), have been identified in filarial nematodes and shown to influence embryogenesis (5), it is not clear if they play any role during the molting process. A number of enzymes including cathepsins, collagenases, lipases, Zn-metalloproteases and aminopeptidases have been implicated in the ecdysis of L3 larvae (6–8). Transcriptional data from microarrays indicated that the transition of the L3 from the vector (at ambient temperature) to the mammalian host (37°C) involves the induction of expression of a wide variety of genes termed adaptation- and infectivity-associated genes (9). While the depletion of the endosymbiotic bacterium *Wolbachia* by antibiotics results in sterility of adult female parasites and disrupts larval molting (10), the significance of this symbiosis, however, in the molting process is not clear as filarial (and other) nematodes that are *Wolbachia*-free also molt quite successfully.

Because this developmental process can be recapitulated readily *in vitro*, in serum-free conditions where the metabolic requirements and signals necessary for the induction of the L3 to L4 molt have been defined (11, 12), we assessed the proteomic profiles of the L3-L4 molt to understand molecularly the initiation of the molting process, and subsequently identify and target specific crucial pathways that could prevent parasite development. In the process, we also defined the protein expression profiles of the endosymbiont, *Wolbachia* (*w*Bm).

## Results

### *B. malayi in vitro* molting protein atlas

The *in vitro* developmental molt of *B. malayi* from the infective L3 larvae to the L4 stage was partitioned into 9 segments (see Materials and Methods, **Figure 1A**) and analyzed by liquid chromatography-tandem mass spectrometry (LC-MS/MS). At a false discovery rate (FDR) of 0.01, a total of 3466 proteins of *B. malayi* (**Table S1**) and 57 of *Wolbachia* (*w*Bm) origin were identified (**Table S2**). Comparative expression profiles of *Brugia*-derived proteins of the quadruplicates across all the time-points using Spearman rank correlations revealed two distinct groups (**Figure 1B**), an early phase (L3, 3Hrs and 24Hrs) and a mid-late phase, the latter being able to be further divided into an early ascorbic or middle phase (5 days to 24HrAsc) and late phase (48HrAsc to L4). Principal component analyses further highlight the changes in protein expression profiles between the ascorbic phases (5 Days, 3HrAsc, 24HrAsc, 48HrAsc), Molting and L4 stages (**Figure 1C**).

**Figure 1.**
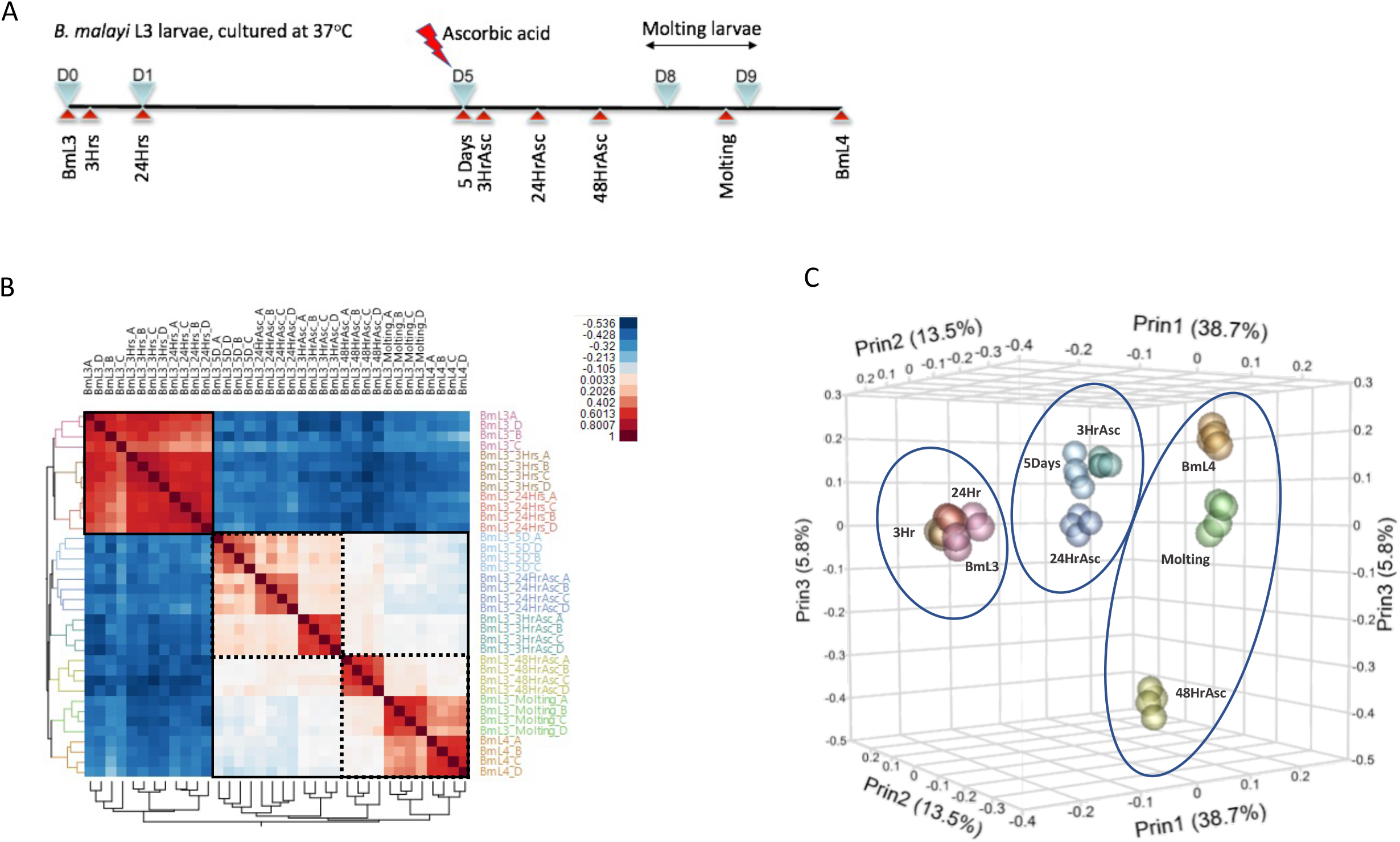
Overview of the B. malayi L3 molting proteome. A) The time line for the L3 to L4 molting, with days (inverted blue triangles) and the time-points profiled (red triangles). B). Correlation heat map of quadruplicates from each time point. Red to blue denotes higher to lower abundance of detected protein. The early and later (middle, late) phases are highlighted with the black squares C). Three-dimensional principal component analyses plot highlighting the proteomic signature profiles across the time points of the molting process

On average ~1500 proteins were identified from each stage by at least one unique peptide (**Table S1**). The proteins were identified either exclusively in one stage, commonly or randomly expressed across the stages, or at specific periods during the developmental molt. The identified proteins were placed into defined clusters using t-SNE dimensionality reduction (13). This approach highlights additional sub-clusters of proteins that in turn define transitions between early-mid, mid-late, common and random expression (**Figure S1-A**). For example, the ‘early’ group comprised of three distinct sub-groups that define the stages (L3, 3Hrs 24Hrs) during which sets of proteins were expressed at high levels. Though the vast majority of proteins were detected in most stages, certain clusters of these common proteins were observed to be present in higher abundance across each of the stages (**Figure S1-B**).

Because the protein expression profile appeared to be broadly set into the early, middle and late phases, the expression data were also visualized using supraHex (14) a self-organizing and visualization tool (**Figure 2**). Protein clusters during early development (3Hrs to 5Days) were shown by increased expression that included cathepsin-L like cysteine proteases (Bm7675, Bm7676, Bm7677, Bm7679, Bm7681), cystatin (Bm366), SCP-like extracellular protein (Bm4233) and conserved secreted proteins (Bm16893, Bm16894, Bm16896) among others (15). The later phases of molting were reflected by increased expression of enzymes and proteins involved in cuticle synthesis. The comparisons of the enriched GO categories of the differentially expressed proteins plotted by semantic similarity (**Figure S2**) between early, middle and late phases highlight early cysteine-type peptidase catalytic activity (in Early <-> Middle; Early <-> Late), oxidoreductase activity, steroid dehydrogenase activity and peroxidase and protein-disulfide oxidoreductase activities (in Middle <-> Late).

**Figure 2.**
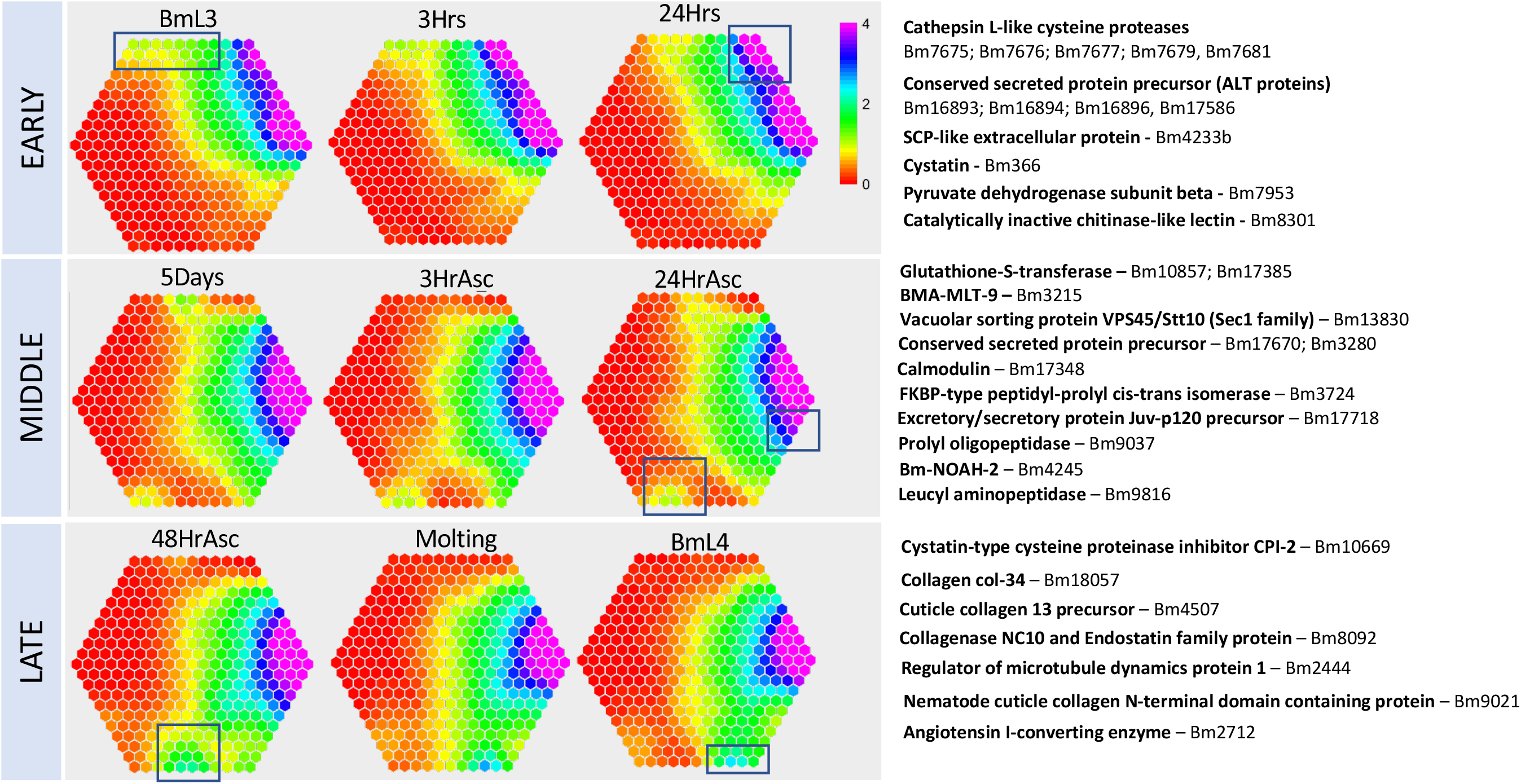
Clusters of molting proteome. The supraHex maps illustrate sample-specific expression profile, where proteins with similar expression profile are mapped to the same position or cell. The nine time-points are broadly placed into three phases (early, middle and late) are shown. The proteins clustered in the highlighted cells are shown to the right. The color bar codes for expression levels [log2(intensity)], from violet to red denoting high to low expression.

### Functional enrichment

Because not all proteins can be classified through GO categories, *B. malayi* proteins identified during the molting process were classified into functional groups (16, 17). The distribution of the functional groups (plotted as percentage of proteins identified from each stage) again appears to be clustered into three broad groups (early, middle and late) (**Figure 3A, Supplemental Figure S3A, B**). Along similar lines, gene set enrichment analysis indicated enrichment for secreted and energy metabolism-related proteins during the early phase (L3, 3Hrs and 24Hrs) compared to the later stages (middle and late). While the energy metabolism-related proteins comprised of dehydrogenases and oxidoreductases, the secreted class of proteins were primarily the cysteine proteases, abundant larval transcripts, serpins, cystatins and several conserved hypothetical proteins.

**Figure 3.**
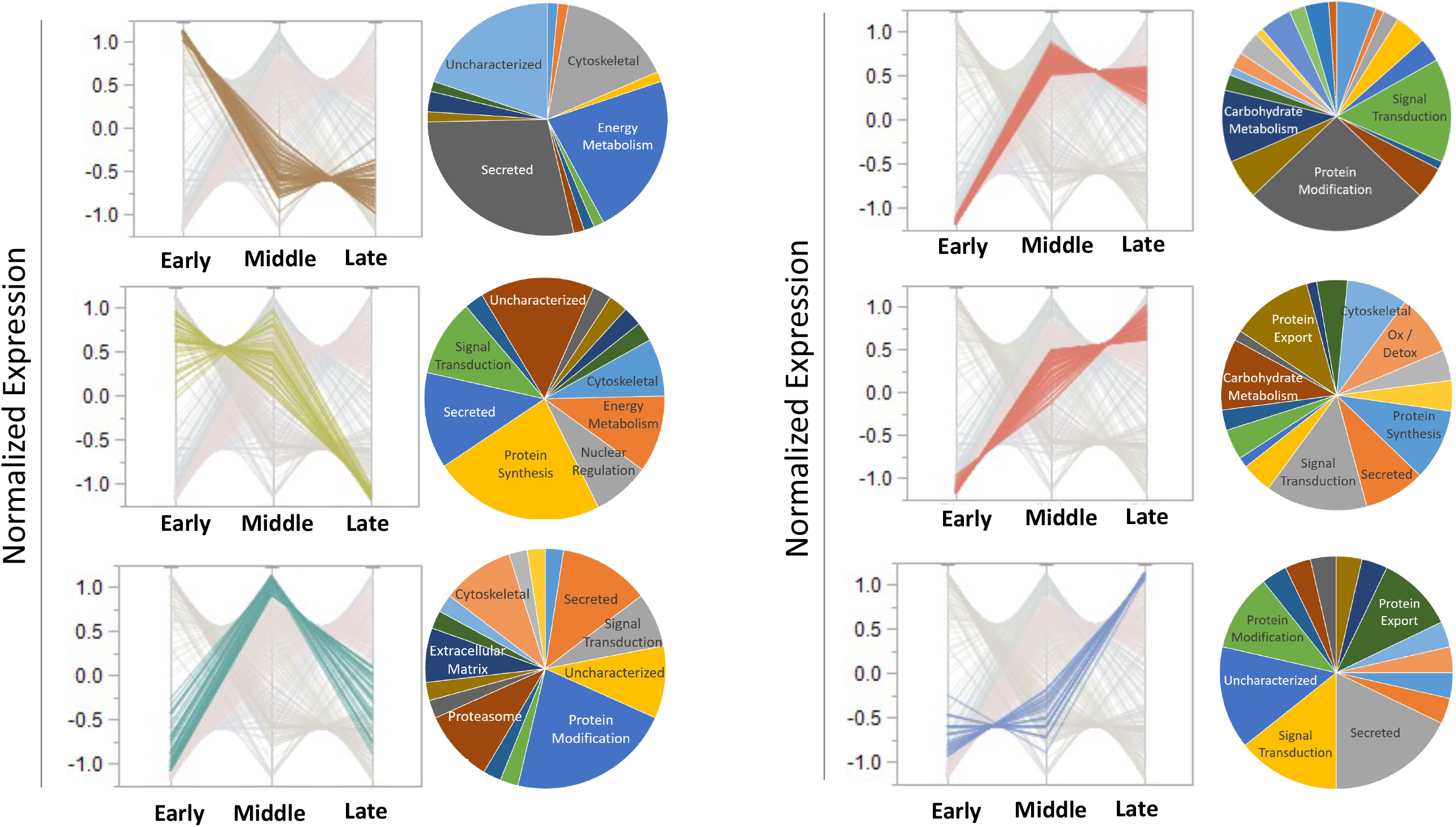
Functional analysis of differentially expressed proteins. The line graphs represent the significantly enriched proteins during the early, middle or late phases, while the pie charts represent their functional classifications. The values on the x-axis denotes the normalized expression levels

In contrast, enrichment (FDR < 0.01) of proteins involved in lipid metabolism, protein export and protein modification were observed during the mid-to-late phase (5days to L4), compared to the early phase (L3, 3Hrs and 24Hrs) (**Supplemental Figure S3B, C**). The addition of ascorbic acid resulted in the enrichment of secreted proteins (**Supplemental Figure S3D**) that were distinct from the secreted proteins during the early phase (**Figure 3**) and annotated as conserved hypothetical proteins and extracellular matrix-related proteins primarily composed of collagens and related machinery.

### Stage-specific expression of cysteine proteinases

Phylogenetic analyses of all the *B. malayi* encoded cysteine proteases indicated that the majority of cathepsin L-like cysteine proteases detected during the L3 to L4 developmental molt were similar to BmCPL-1, BmCPL-4 and BmCPL-5 (**Figure 4A**), and very much similar to what has been observed in *O. volvulus* (18)*, B. pahangi* (19) and *D. immitis* (20). Cathepsin A (Bm3985, Bm6297), Cathepsin B (Bm2365), and Cathepsin F (Bm1575, Bm3996) were increased during the ascorbic acid phase. Interestingly, compared to the enrichment of CPL-1, -4 and -5 observed in the L3 to L4 stages, combined analysis with the protein profiles of the stage-specific somatic proteomes (16) and transcriptomes (21) indicated the stage-specific expression and/or utilization of Bm12799, Bm12798, Bm12797, Bm8207, Bm8172, Bm748 and Bm99 (CPL-2, −3, −6 and −7 like) by the immature and mature stages of microfilariae (**Figure 4B**).

**Figure 4.**
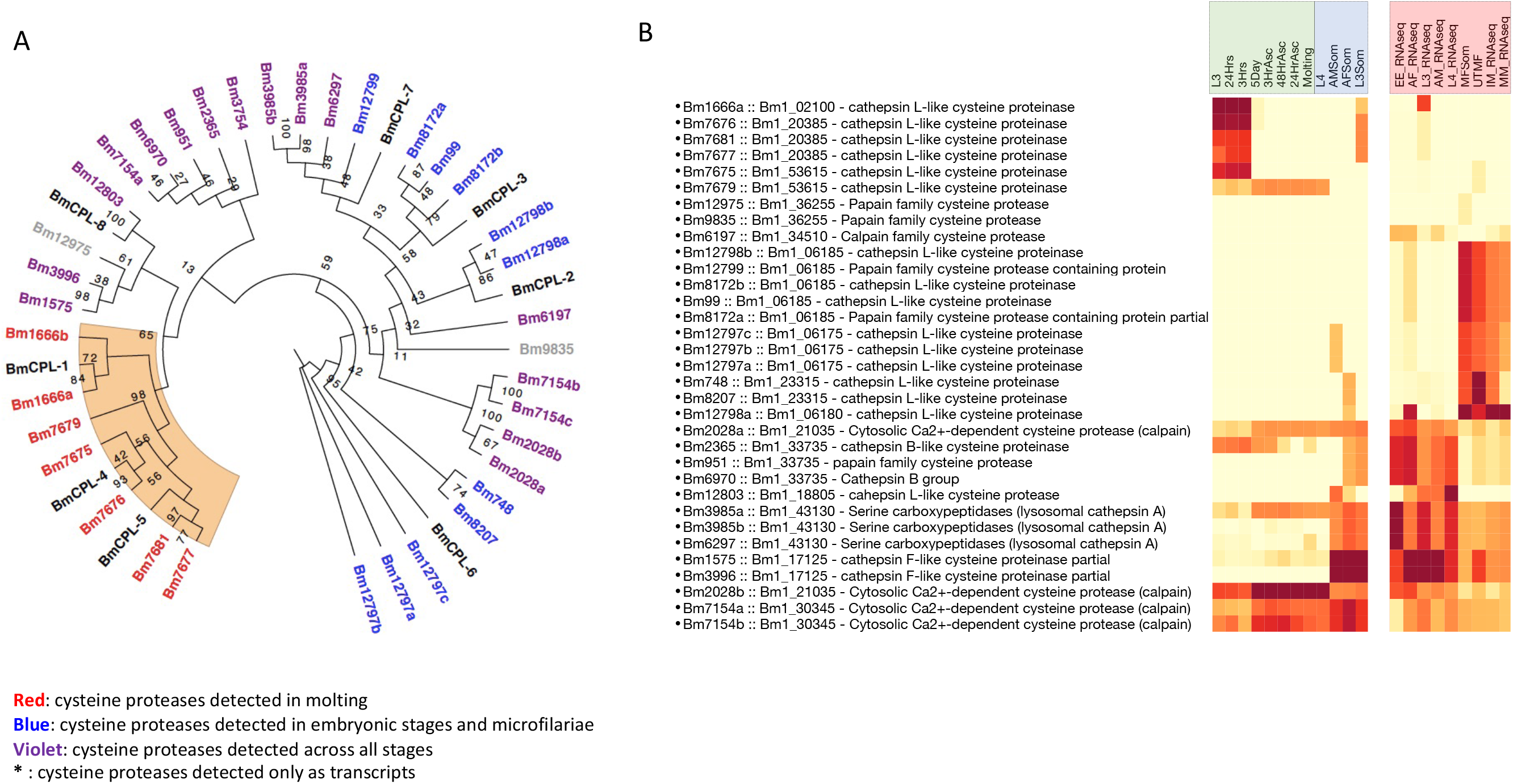
Cysteine proteases of *B. malayi*. A). Maximum likelihood phylogenetic analyses of the *B. malayi* cysteine proteases using RaxML under the conditions of the GAMMA model with nodal support values generated through 1000 bootstrap replicates. The tree depicts the proteases detected during the molting process (red), are primarily BmCPL-1, −4 and −5 like cysteine proteases. The proteases with previous proteomic and/or transcriptomic evidence in embryonic stages and microfilaria (blue) or constitutively across all stages (violet), and known *B. malayi* cysteine proteases (black, prefixed with BmCPL). B) Heatmap depicts the expression and clustering of the cysteine proteases detected in the current study (green), with the stage-specific somatic proteomes (blue) or stage-specific transcriptomes (Pink). The corresponding PubLocus annotation is also provided as the somatic proteome and transcriptomes were based on the previous annotation.

Since the cathepsin’s have been hypothesized to be stored in the granules of the glandular esophagus and transported through pseudocoelomic fluid networks and secretory vessels to the hypodermis during molting(18), the cathepsin activity was localized using ProSense 680, a fluorescent catabolic substrate of cathepsin at 24hrs and 5days. As shown in **Figure 5A** cathepsin activity was more pronounced near the hypodermal and sub-cuticular areas at day 5 compared to that seen in the area around the gut/pharynx at 24hrs. As observed previously with other filarial parasites, inhibition studies of cathepsins with chemical inhibitors were carried out to assess the impact of cathepsins in this *in-vitro* molting model (19, 22). Similar to the studies with other filarial parasites, a dose-dependent inhibition of molting with Z-Phe-Ala-FMK was observed (**Figure 5B**). In addition, a novel cysteine protease inhibitor - K11777 was much more effective than Z-Phe-Ala-FMK, and at concentrations above 2 μM, blocked the development of L3 larvae *in vitro* and completely killed all the larvae within 24hrs (**Figure 5C**). At 1 μM K11777 larvae were still viable at 4 days but failed to molt.

**Figure 5.**
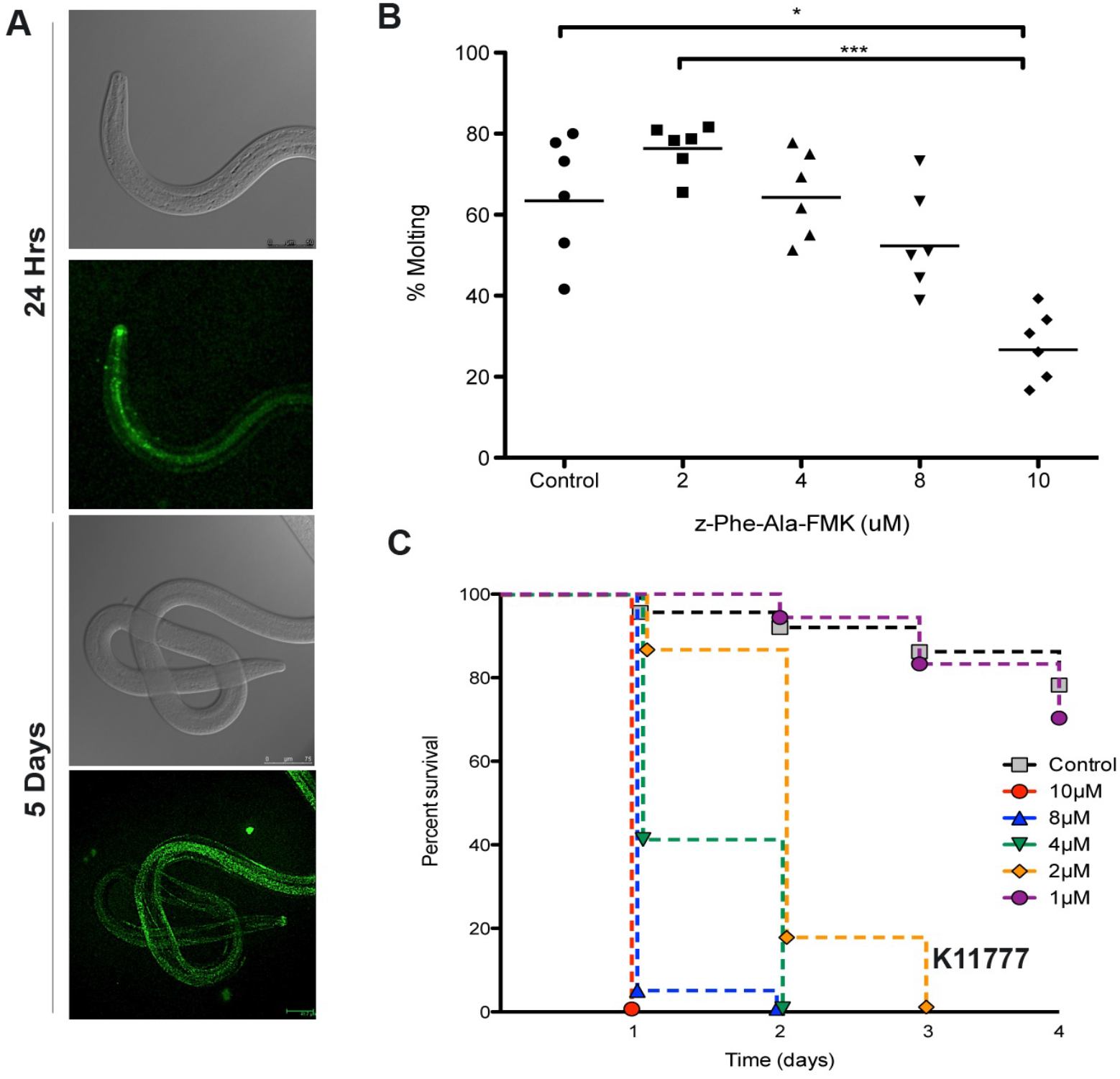
Cathepsin Activity. A) Confocal images showing the breakdown of ProSense680 fluorescent substrate in the gut/pharyngeal tract at 24hrs and hypodermal areas by day 5; B. Inhibition of cathepsin activity with z-Phe-Ala-FMK shows a dose dependent (2 μM to 10 μM) inhibition of molting in L3 larvae at 10 days. C. Inhibition and death of L3 larvae by the cathepsin inhibitor K11777, plotted as survival graph over 4 days. Each dot/time point represents the average of 10 wells with ~10 larvae in each well.

### Kinases

Ascorbic acid is known to regulate a wide variety of biochemical processes, of which the activation/inhibition of kinases and collagens are most notable (23–26). To understand the effect of ascorbic acid in inducing the crucial developmental molt, the expression of the *B. malayi* kinome across the L3-L4 development was analyzed. A total of 426 eukaryotic protein kinases (ePK) were identified, comprising 2.5% of the predicted proteome of *B. malayi*. It appears that *B. malayi* may be missing the core kinases CAMK/RAD53 and CMGC/RCK/MAK. Interestingly, the predominant clustering of the kinases and their relative abundance upon stimulation with ascorbic acid appears to activate and induce the expression of CAMK and AGC family of kinases (**Figure 6**), that are known to regulate cytoskeletal reorganization and extracellular matrix remodeling, and are potential therapeutic targets (27, 28).

**Figure 6.**
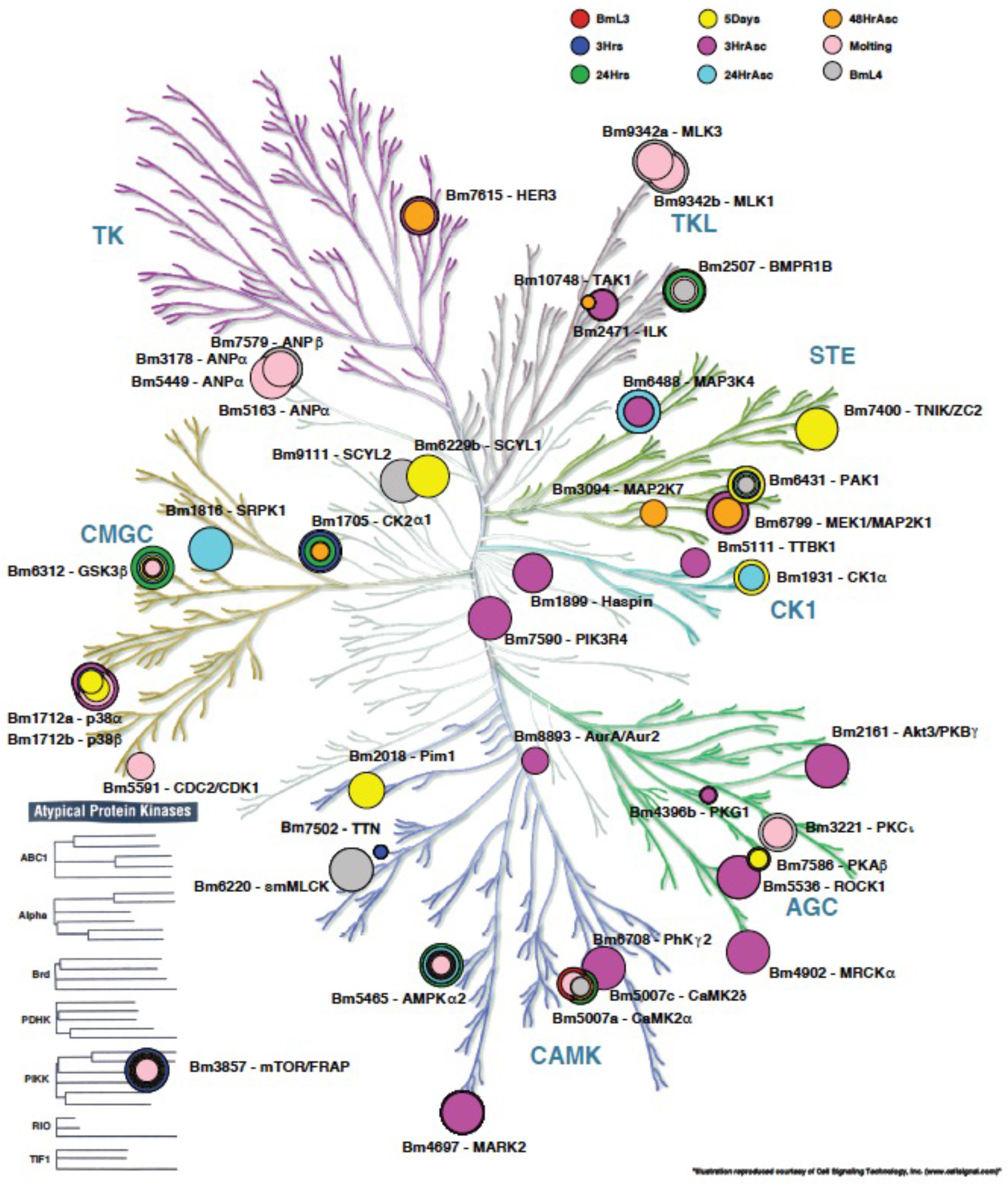
The kinome of *Brugia malayi* during molting. The figure denotes the kinases (annotated with *B. malayi* accession and kinase name) detected across the various time points during the molting process overlaid on the phylogenetic tree of kinases with the major kinase family groups (CMGC, AGC, CAMK, STE, TKL, TK, atypical protein kinases). The size of the circle denotes the intensity of expression. Concentric rings denote expression in multiple stages at varying intensities.

### Cuticle, Collagens and Collagen-machinery

Clustering of representative members of *C. elegans* collagens (29) with *B. malayi* collagens detected during the molting process highlighted the specific expression of the *dpy-2*, *dpy-10*, *dpy-7* and *dpy-8* and select Group-2 collagens during the middle phase (**Figure 7A, B**). The clustering data also indicate that constitutively expressed collagens primarily belong to Group-1 collagens (**Figure 7B**).

**Figure 7.**
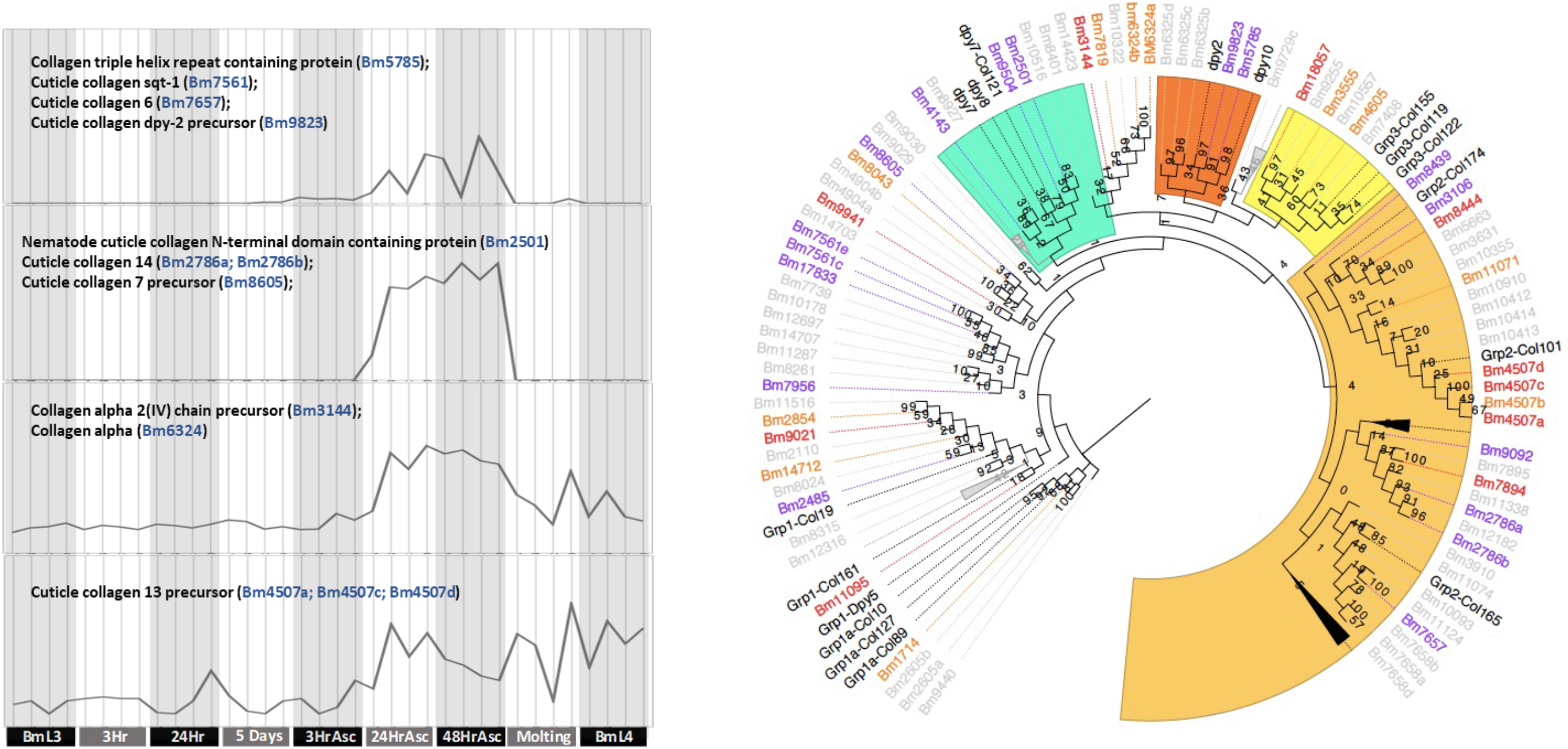
Collagens and cuticle machinery. A) Parallel plots depicting clusters of enhanced expression of collagen proteins upon induction with ascorbic acid during the molting process. The vertical lines denote the quadruplicates at each time point B) Phylogenetic tree based on *C. elegans* collagen groups (black), indicates preferential utilization of group2 and ‘dpy’ like collagens during the ascorbic phase (violet), and a more distributed constitutively expressed (orange) or during late phase (red). Collagen proteins not detected are in grey.

The synthesis and trimerization of collagen fibers is a series of complex enzymatic processes involving prolyl-4 hydroxylases, disulfide isomerases, peptidyl-prolyl cis-trans isomerase, blisterases and aminopeptidases (30). The clustering of the collagen machinery proteins identified three well-defined clusters (**Figure S4**). Though the exact combination of molecules involved in this process is unknown, it is likely that collagen machinery components upregulated at 5-days and beyond are responsible for the generation of the new cuticle. Surprisingly, even though nuclear hormone receptors are known to play a role in molting, there was barely any expression of the nuclear hormone receptors of *B. malayi* identified at the protein level.

## Discussion

To establish a filarial infection in a vertebrate host, the infective L3 larvae must undergo a series of successful developmental molts in the host to become adult parasites. The *B. malayi* L3 to L4 molting in a controlled *in vitro* environment devoid of serum or growth factors appears to be the best way to investigate the essential pathway for the early development of the mammalian stage parasites. It should be noted that the L3 larvae had to be shipped post harvesting and hence the proteomic profile might be slightly different if they were to be cultured immediately upon harvest. Although gene expression data of developmental stages in *C. elegans* (31, 32) and microarray analyses of *B. malayi* L3 larvae (9) are available, direct comparative analyses of the *in vitro* molting model data were limited, due to the variable durations of molting processes (relatively short developmental time in *C. elegans* (~9 hours) compared to *B. malayi* (~9 days) and the fact that unlike *C. elegans*, the L3’s of *B. malayi* cannot be synchronized. Further, microarray analyses of *B. malayi* L3 larvae (9), were also not directly comparable because of experimental design constraints. It is also to be noted that all the transcriptomic and proteomics analyses are based on the quality of the annotated genomes available.

The early phase of the L3 developmental process showed that cysteine proteases play an important role during molting, a finding with parallels to studies in *O. volvulus* (33). While the proteomic findings corroborate previous RNA expression data of CPL-1, −4 and −5 (Group Ia) in the L3 stages, it was interesting to note that the cysteine proteases (CPL-2, −3, −6, −7 and −8) that belong to group Ic (19) were most abundant in the microfilarial and intra-uterine microfilarial stages. It has been hypothesized that the CPLs in the L3s are stored in the granules of the glandular esophagus that are transported during molting through pseudocoelomic fluid networks and secretory vessels to the hypodermis and the cuticle (18). Fluorescent substrate catalysis imaging demonstrated cathepsin-like activity in the granules of the esophageal tube as early as 24 hours at 37°C and that remains visible in the hypodermal and sub-cuticular regions of the worm following ascorbic acid induction of molting *in vitro*.

Cystatins, are endogenous cysteine protease inhibitors and when transported to the cuticles of the filarial L3 and L4 larvae during molting (18, 34), they possibly regulate the cysteine protease activity that does not inhibit the formation of the new cuticle, but rather support the separation of the L3 and L4 cuticles *in vitro* (18). The ability to inhibit molting and/or development of *B. malayi* L3 larvae by chemical inhibitors of cathepsins (Z-Phe-Ala-FMK & K11777) demonstrate the critical role these molecules play in the development of the nematode parasite. K11777, a novel cysteine protease inhibitor has been shown to be effective against *Trypanosoma cruzi, Leishmania tropica* and *Schistosoma mansoni* (35). However, although inhibition studies by RNAi have previously been used with *B. malayi* (22, 36–38) and *O.volvulus* (33), *B. malayi* L3 larvae were quite sensitive to dsRNA, siRNA, shRNA targeting the cysteine proteases, as the induction of molting with ascorbic acid in the presence of even non-specific silencing nucleic acids resulted in death of all larvae. Likewise, although electroporation has been used successfully in adult worms (37), it was lethal to *B. malayi* L3 larvae.

The middle and late phases of molting (Day 5 and beyond) were primarily comprised of enzymes and proteins related to cuticle formation. Among the enzymes were protein kinases that are known to be regulated by the diverse biochemical functions of ascorbate and dehydroascorbate and influence the cytoskeletal reorganization. For example, ROK1, MRCKa, AKT3/PKBa facilitate extracellular matrix (ECM) remodeling, while the NIMA-related kinases NEKL-2/NEK8/NEK9 and NEKL-3/NEK6/NEK7, together with their ankyrin repeat partners, MLT-2/ANKS6, MLT-3/ANKS3, and MLT-4/INVS, are essential for normal molting (25). The NIMA-related kinase network functionally interacts with CDC-42 and SID-3/ACK1 to facilitate ECM remodeling. It would be interesting to investigate further the role of CAML and AGC family of kinases expressed post ascorbic acid in the molting process. Ascorbic acid also promotes the expression of procollagen C-proteinase enhancer (PCPE-1) (39), necessary for collagen fibril assembly. Though the timing was not as expected, the *B. malayi* orthologue of PCPE (Bm3917) was detected at 48hr post-ascorbic acid.

Nematodes do have an ascorbate biosynthetic pathway that is significantly different from that found in animals, plants and fungi (40). Although, the homologues of the mammalian nucleobase ascorbate transporter or nucleobase cation symport 2 (NAT/NCS2; SCVT1 and SCVT2) (41) that support the active transport are not present in *B. malayi,* the conserved NAT motif [QEP]NXGXXXXT[RKG] could be mapped to Bm2885. However, Bm2885 protein was not detected in any of the stages.

Temporal regulation of the collagen gene expression in *C. elegans* has been shown to occur during the molting period (42, 43). The expression patterns of various classes of collagens were expressed at varying levels across the various stages or specifically enriched in stage-specific molts (44, 45). Though the cuticles of larval and adult stages appear to be similar both structurally and biochemically (46–48), the preferential utilization of collagens by L3 larvae that cluster with Group 2 collagens (29) corroborates data previously described (49, 50). The production of collagen influenced by ascorbate in humans has been largely attributed (primarily as a co-factor) to prolyl hydroxylation (51) and/or increases in steady-state levels of procollagen mRNA (52). However, other studies suggested that the role of ascorbate in collagen synthesis may be unrelated to hydroxylation (53). In addition, there have been other studies that have suggested that this process may reflect a lack of transcriptional regulation (54) or may involve the protein synthesis machinery or other mechanisms (38, 55). Complementation of *B. malayi* encoded collagen enzymatic machinery genes in *C. elegans* suggested both conserved and divergent functional activities (reviewed in (30)).

The association between the filarial parasites and their endosymbiont *Wolbachia* (*wBm*) is ancient (in evolutionary terms) with mutual, symbiotic interrelationships (56). Data from studies targeting *Wolbachia* with antibiotics not only supports this symbiotic relationship, but they also highlight the dependence of the filariae on the bacteria for a diverse range of biological and stage-specific processes (57). Although *B. malayi* does not have a complete gene set required for the *de novo* purine synthesis and may be dependent on *wBm*, we were unable to observe this directly for the reason that the culture medium was supplemented with ribonucleosides and deoxyribonucleosides. Likewise, the influence of *Wolbachia* in the molting process was not clear in the molting process. Although a total of 57 Wolbachial proteins were detectable, there was no obvious phase-specific discernible patterns (**Supplementary Figure S5**), except for Chaperonin GroEL (HSP60; Wbm0350) that was detected during the molting and L4 stages only. Only 4 Wolbachia proteins (Wbm0495-molecular chaperone DnaK, Wbm0793 – Type IV secretory pathway VirB6 components, Wbm0635 – RecG-like helicase, Wbm0786 – valyl-tRNA synthetase) were detected across each of the time points. Though non-*Wolbachia* containing filarial parasites (e*.g. Loa loa*), having similar genomic structures to those of *B. malayi* and *W. bancrofti* (58) and have normal L3-L4 molts, indicates no active role for *Wolbachia* in the molting process. It is likely, however, that the low coverage of *Wolbachia*-derived peptides was influenced by the limitations associated with mass spectrometry and highly limited *wBm* numbers and protein content during the L3-L4 transition. *Wolbachia* can be depleted by antibiotics belonging to the tetracycline class of antibiotics and causes sterilization of the adult females and blocks development of the parasites. Because the *in vitro* molting efficacy was significantly inhibited by tetracycline, and by a chemically modified tetracycline (that lacks anti-microbial activity) in the absence of *Wolbachia* clearance suggested disruption of filarial physiology as a possible mechanism (59, 60).

In conclusion, using a defined *in-vitro* model of filarial molting that limits the possibility of any extraneous or unknown mediators influencing the molting process our data highlights the minimal set of proteins and processes needed for the *B. malayi* L3 larvae to molt to L4. Moreover, a number of these are very good targets for prophylactic and/or therapeutic intervention, in other non-human filarial parasites.

## MATERIALS AND METHODS

### Parasites and molting model

*B. malayi* L3 larvae were obtained under contract from the FR3 facility at University of Georgia to the NIAID. The L3 larvae were cultured (**Figure 1A**) as described previously (61). The larvae were snap frozen immediately upon processing (‘BmL3’; D0), post-incubation at 37°C for 3 hrs (‘3Hrs’), 24 hrs (‘24Hrs’; D1), 5 days (‘5Days’; D5). Following the addition of ascorbic acid (75 μM) at day 5, parasites were collected 3 hrs later (‘3HrAsc’), 24 hrs later (‘24HrAsc’), and 48 hrs later (‘48HrAsc’) and then again during molting (‘Molting’; Days 8/9) and finally as viable L4 larvae (‘BmL4’). The animal procedures used for the life-cycle of *B. malayi* were conducted in accordance with the animal care and use committee guidelines at the National Institutes of Health and at the University of Georgia.

### Protein extraction and Mass Spectrometry

Total soluble proteins were extracted from ~500 larvae from each time point using the UPX universal protein extraction kit (Protein Discovery, San Diego, CA) as per the manufacturer’s instructions. The protein concentrations were estimated using Pierce BCA protein assay kits. The protein samples from L3 to L4 larvae stage were reduced, alkylated and trypsin digested overnight following filter-aided digestion procedure using a FASP digestion kit (Protein Discovery, San Diego, CA). Tryptic peptides were further desalted using C18 spin columns (Thermo Fisher Scientific, IL). Samples were then lyophilized and reconstituted in 0.1% trifluoroacetic acid to be analyzed in quadruplicates without fractionation for quantitation purposes.

Nanobore RPLC-MSMS was performed using an Agilent 1200 nanoflow LC system coupled online with LTQ Orbitrap Velos mass spectrometer. The RPLC column (75 μm i.d. x 10cm) were slurry-packed in-house with 5 μm, 300Å pore size C-18 stationary phase into fused silica capillaries with a flame pulled tip. After sample injection, the column was washed for 20 min with 98% mobile phase A (0.1% formic acid in water) at 0.5 μl/min. Peptides were eluted using a linear gradient of 2% mobile phase B (0.1% formic acid in ACN) to 35% B in 100 minutes, then to 80% B over an additional 20 minutes. The column flow-rate was maintained at 0.25 μl/min throughout the separation gradient. The mass spectrometer was operated in a data-dependent mode in which each full MS scan was followed by sixteen MS/MS scans wherein the sixteen most abundant molecular ions were dynamically selected for collision-induced dissociation (CID) using a normalized collision energy of 35%.

The LC-MS/MS data were processed using PEAKS 7 Studio (ver 7, Bioinformatics Solutions Inc.). MS/MS data were searched against a combined decoy database of *B. malayi* (http://parasite.wormbase.org; WBPS10) and *Wolbachia* (*wBm*) containing both forward and reverse sequences as well as a common contaminant database using default parameters. Dynamic modifications of methionine oxidation and N-terminal acetylation as well as fixed modification of carbamidomethyl cysteine were included in the database search. Only tryptic peptides with up to two missed cleavage sites with a minimum peptide length of six amino acids were allowed. The false discovery rate (FDR) was set to 0.01 and threshold-based filtering of −10logP scores of 30 for both peptide and protein identifications. Statistical tests were performed on normalized spectral abundance (NSAF, relative abundance) of the proteins to determine significant protein changes between time points (**Table S1**). Rarefaction analyses were carried out using the number of peptides identified in each sample (Supplemental Figure S6). The data were analyzed using JMP Genomics 9.0 (SAS Institute Inc) and R (3.6).

### Differential Analysis

Gene Set Enrichment Analysis (GSEA, Broad Institute), a method for analyzing molecular profiling data, examines the clustering of a pre-defined group of genes or proteins (gene set) across the entire database in order to determine whether the gene set has biased expression in one condition (or stage) versus another (62). For this analysis, the entire list of *B. malayi* L3 larval molting-associated proteins were sorted on their relative abundance. The distribution of proteins from an *a priori* defined set throughout this ranked list was then determined using GSEA (as described previously (63)). Sets of genes encoding for proteins in each functional category were analyzed using GSEA for specific enrichment of genes/proteins.

Differential expression of proteins between early, middle and late phases of the molting process and also the early, middle or late phases with the other two phases were analyzed by QPROT(64), an extension of QSPEC(65) suite for label free proteomics. Gene Ontology enrichment for the differentially expressed proteins was analyzed using R implementation of TopGO (66) and semantic similarity graphed using REVIGO (67).

### Cathepsin Activity and Chemical inhibition

Prosense 680 Fluorescent Imaging Agent (Perkin Elmer #NEV10003, previously VISEN) was used to visualize the activity of cathepsin’s *in vivo* after incubating the larvae for various time-points at 37°C. Fluorescence images were collected on a Leica SP5 X-WLL confocal microscope (Leica Microsystems, Exton, PA). Z-Phe-Ala-FMK (Sigma # C1480) was prepared as 10 μM stocks in DMSO. L3 larvae were cultured in the presence of 2 μM, 4 μM, 8 μM and 10 μM of Z-Phe-Ala-FMK in L3 media for 5 days. The numbers of shed cuticles (as indicator of successful L3 larval molt) were enumerated on Day 10. K11777, a novel drug that targets cysteine proteases (35) was also tested at concentrations of 2 μM – 10 μM.

### Kinome analyses

The kinome of *B. malayi* was analyzed using Kinannote (68) and the intensities (expression abundance) of the kinases was plotted using a slightly modified version of Kinomerender (69).

### Sequence analysis

Sequences of the cysteine proteases, collagens (*B. malayi* and *C. elegans*) were aligned in MegAlign Pro (DNASTAR, Lasergene 17) using MUSCLE algorithm. Next, maximum likelihood (ML) trees and bootstrap (BS) trees were generated with a final “best” tree generated from the best scoring ML and BS trees using RAxML v8.2.10. The resulting phylogenetic trees were visualized in FigTree 1.4 (http://tree.bio.ed.ac.uk).

## Acknowledgements

This work was funded in part by the Division of Intramural Research, National Institute of Allergy and Infectious Diseases, National Institutes of Health.

---

**Figure S1.**
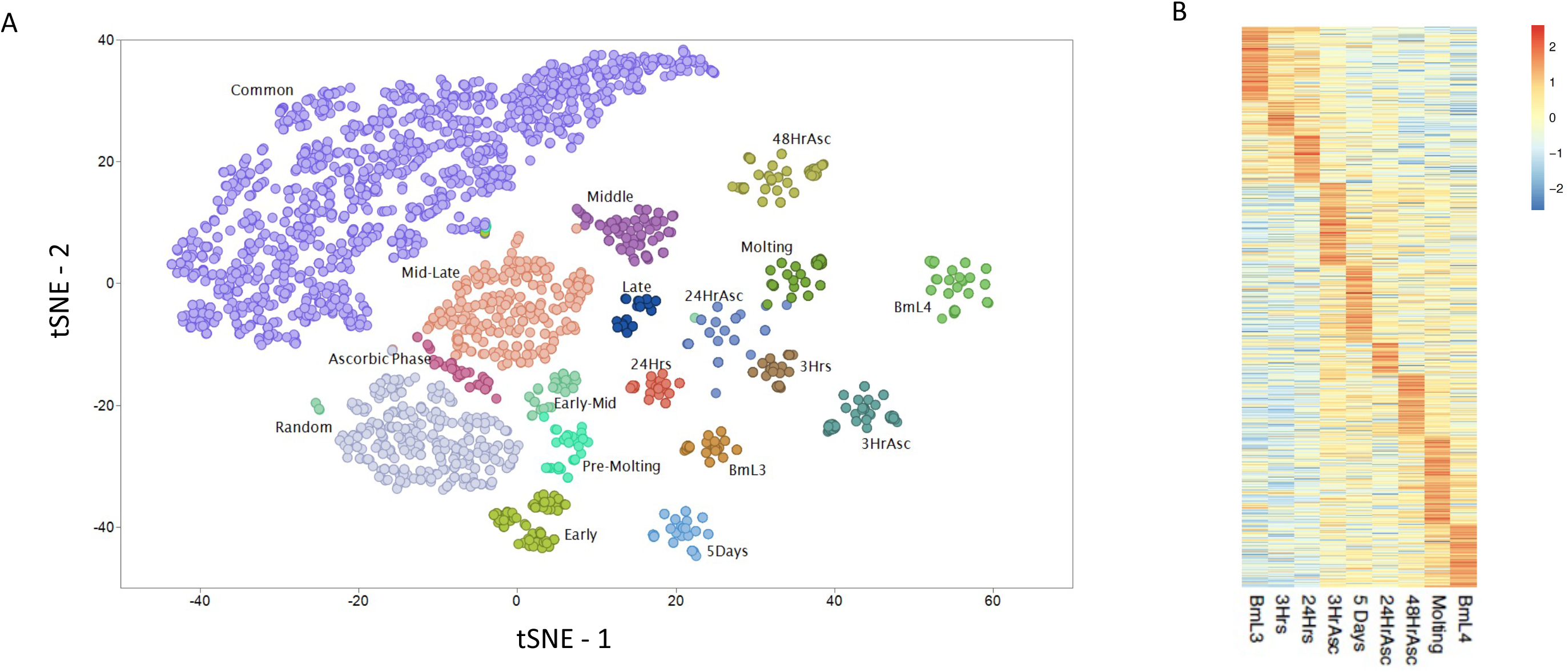
Stage-specific proteomic expression. A) Two-dimensional t-SNE plot of the protein abundance. Each point represents a protein colored by the cluster group B) Circular polar histogram of protein abundance across the clusters defined by t-SNE. The abbreviated pie names: EM – Early to Mid; L – Late; PM – Pre-Molting; Asc – Ascorbic Phase; Asc1 – 3HrAsc; Asc2 – 24HrAsc; Asc3 – 48HrAsc; Molt – Molting. The concentric circles represent the stage-specific proportion of the protein. C) Heatmap of the proteins defined as ‘common’ by t-SNE, highlighting the increased abundance in specific time-points. D) Heatmap of the proteins detected only during early, middle and late phases. Red to blue denoted high to low abundance for both the heatmaps.

**Figure S2.**
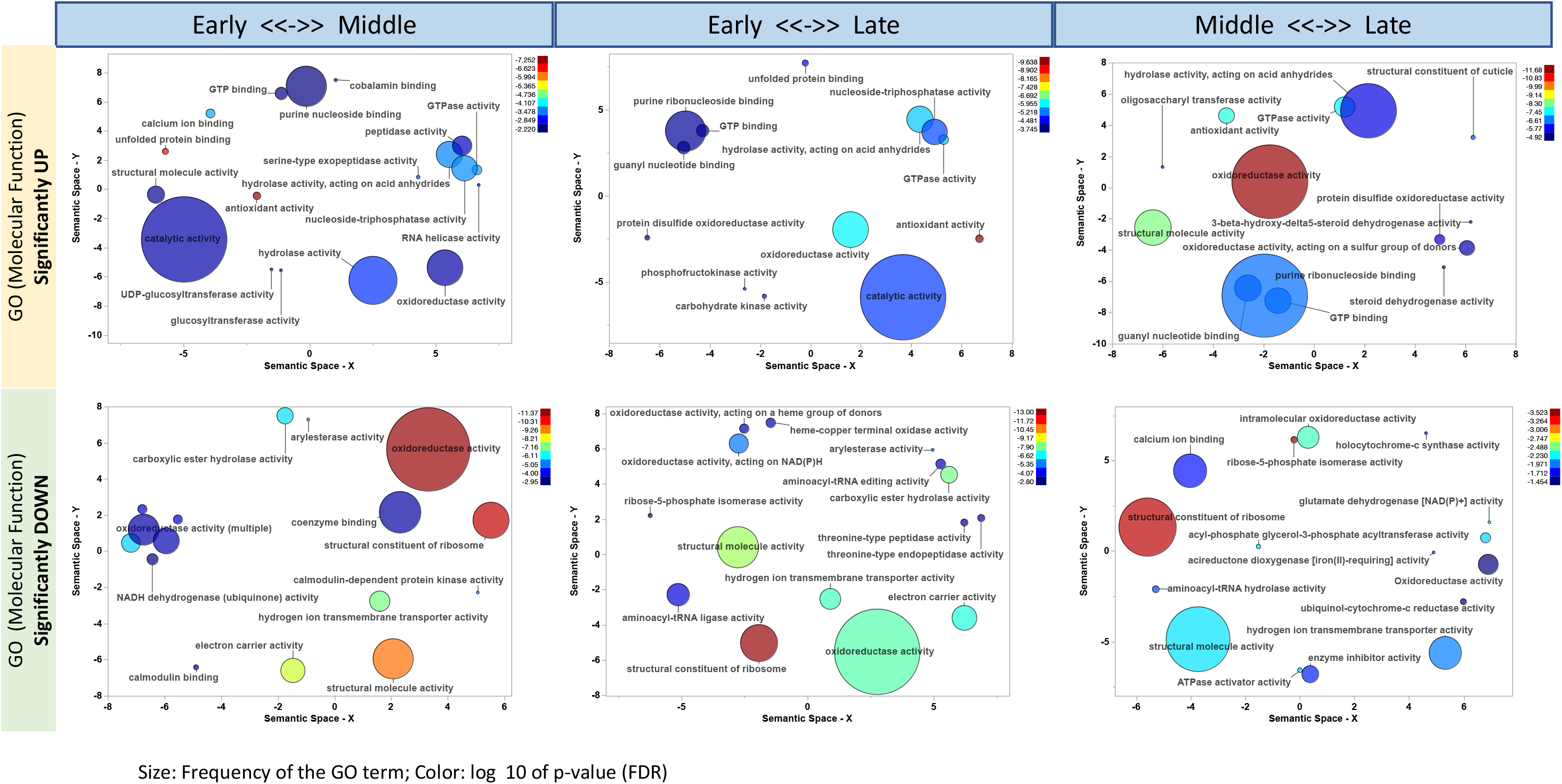
GO enrichment. The scatterplot shows the cluster representatives (i.e. terms remaining after the redundancy reduction) in a two-dimensional space derived by applying multidimensional scaling to a matrix of the GO terms' semantic similarities. Bubble color indicates the user-provided p-value (legend in upper right-hand corner); size indicates the frequency of the GO term in the underlying GOA database (bubbles of more general terms are larger).

**Figure S3.**
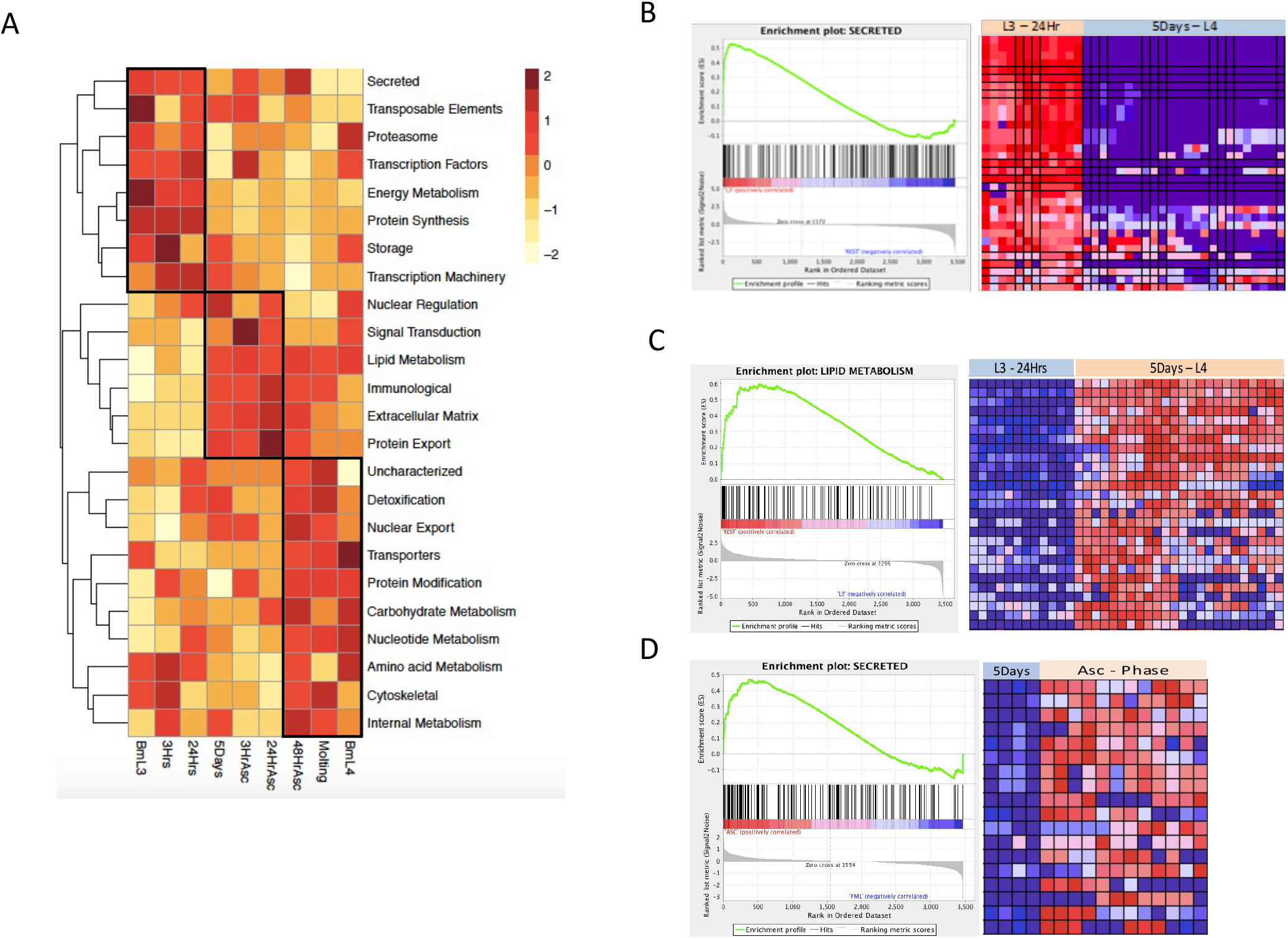
Functional classification and enrichment. A) The heatmap depicts the number of proteins classified into functional groups. Red to yellow denotes high to low. B-D) The gene set enrichment graphs depicting the enriched state of secreted proteins during early phase (B), lipid metabolism during the mid-late phase (C), and distinct group of secreted proteins during the ascorbic acid phase (D). Left half of GSEA plots shows the enrichment score with heat maps of the corresponding proteins from quadruplicates from each set on the right. L3-24hrs set comprises of Fresh L3, 3Hr and 24Hr of culture at 37°C; 5Days-L4 set includes 5days, 3HrAsc, 24HrAsc and 48HrAsc, molting and L4; Asc-Phase includes 3HrAsc, 24HrAsc and 48HrAsc. Red to blue indicates higher to lower expression.

**Figure S4.**
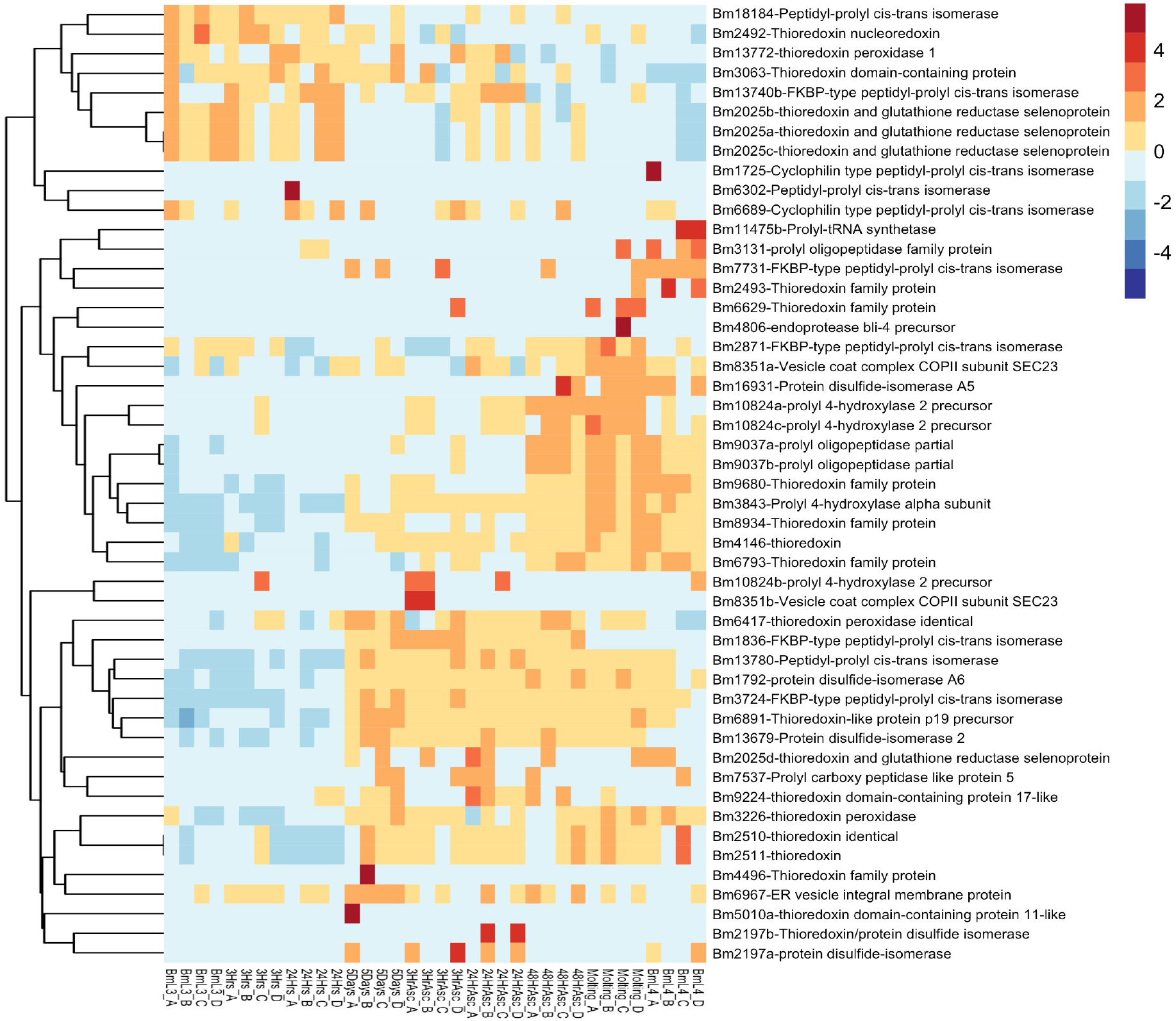
Collagens and cuticle machinery. Heatmap depicts the clustering of proteins involved in the cuticle synthesis. Red to blue denotes high to low expression.

**Figure S5.**
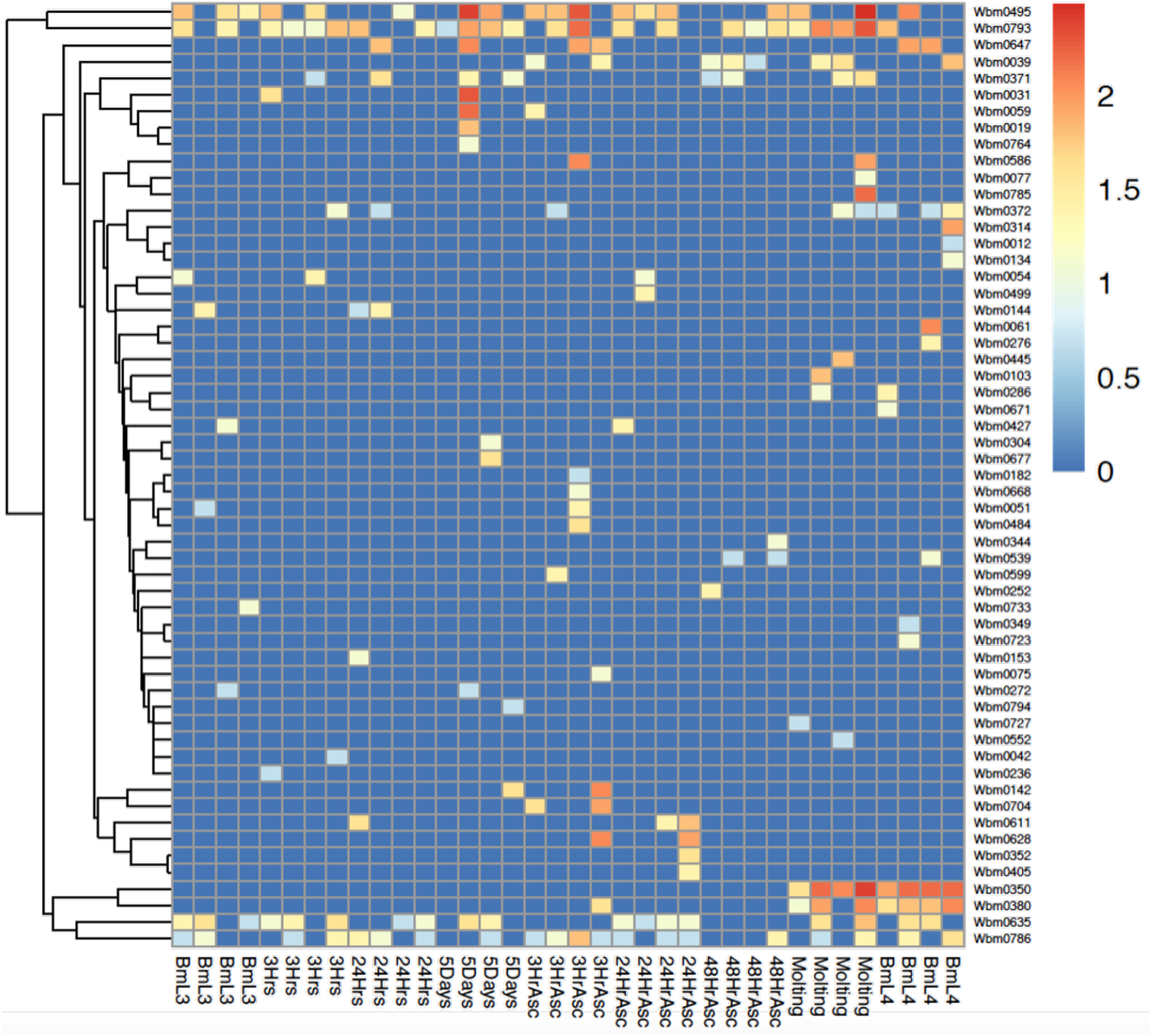
Wolbachial (*wBm*) proteins. The heatmap shows the wolbachial proteins identified during the L3-L4 developmental molt. Red to blue denotes high to low expression.

**Figure S6.**
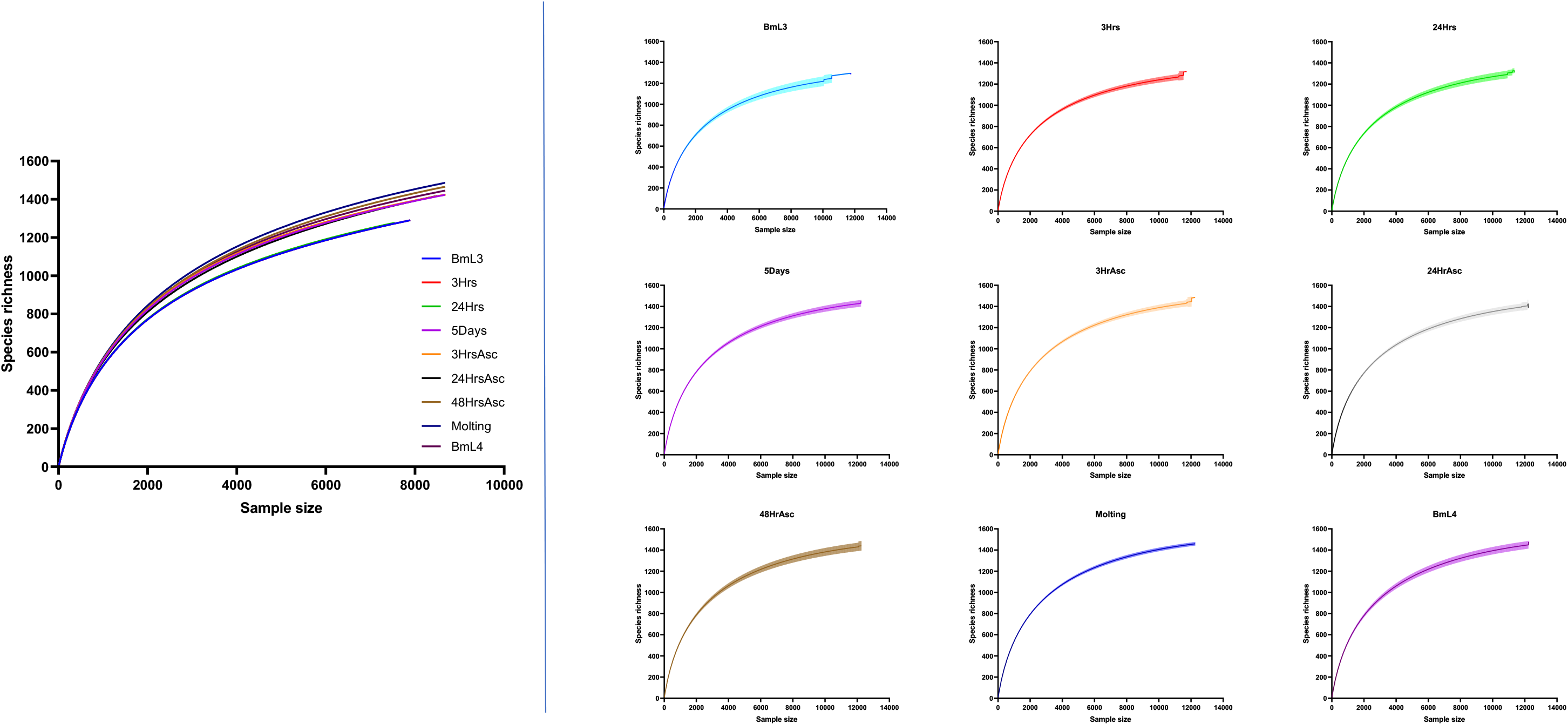
Rarefaction Curves. Rarefaction curves as function of number of peptides identified across the time-points (left panel), and the replicates within each time-point (right).

**Table S1. Normalized spectral abundances of *B. malayi* proteins**

The table lists the normalized spectral abundances of *B. malayi* proteins identified during each of the time-points of the molting process. A, B, C and D represent quadruplicates from each time-point. The significant changes observed between the early, middle and late phases are represented by the log fold changes, false discovery rate (FDR), signal to noise (STN) and the corresponding p-values. The functional classification is also listed.

**Table S2. Normalized spectral abundances of *Wolbachia* proteins**

The table lists the normalized spectral abundances of Wolbachia proteins identified during each of the time points of the molting process. A, B, C and D represent quadruplicates from each time-point. The functional classification is also listed.

